# The Dynamic Monitoring of Tourism Environment of Emeishan Mountain Based on High Score Remote Sensing Iimage and Image Fusion*

**DOI:** 10.1101/2025.07.24.666543

**Authors:** Yongxing Sun, Jianing Gao, Jiarong Li, Yanqiong Chen

## Abstract

Sustainable tourism is widely advocated as a low-impact development strategy that promotes economic growth while preserving ecological integrity. However, empirical evidence increasingly reveals complex environmental trade-offs associated with tourism expansion, particularly in ecologically sensitive heritage sites. Mount Emeishan, a UNESCO World Cultural and Natural Heritage Site in China, exemplifies a destination where tourism has driven regional socio-economic development through culturally meaningful product innovation. Yet, this growth has also imposed significant ecological pressures, particularly in waste management, where hazardous waste accumulation correlates strongly with increasing visitor numbers—posing a growing threat to local ecosystem stability.

To address these challenges, this study proposes a dynamic environmental monitoring framework integrating high-resolution remote sensing imagery (HR-RSI) and advanced image fusion techniques (IFT). The primary objective is to develop a systematic, data-driven methodology for assessing and mitigating tourism-induced environmental degradation, thereby supporting sustainable management of the Emeishan National Park. Empirical evaluation of the framework demonstrated a system stability rate of 88%, representing a 12.7% statistically significant improvement (p<0.05) over traditional monitoring systems under a sample size of 100 test cases.

Through the synthesis of multi-source remote sensing data, the study quantitatively assessed the impacts of tourism activities on key environmental indicators, including vegetation cover, land use dynamics, and water quality. Results revealed a noticeable decline in vegetation coverage and significant land use transitions, particularly in high-traffic zones. Furthermore, elevated water pollution risks were observed during peak tourist seasons, underscoring the seasonal nature of tourism-related environmental stressors.

These findings provide robust scientific support for policy-makers and tourism authorities in formulating evidence-based environmental protection strategies and adaptive resource management policies. The integration of HR-RSI and IFT offers a scalable and replicable approach for monitoring environmental impacts in heritage-rich, tourism-intensive regions, contributing to the global discourse on sustainable tourism and ecological resilience.

## 1. Introduction

Sustainable tourism has long been characterized as a “green industry” due to its traditionally assumed minimal environmental footprint. However, a growing body of empirical research reveals increasingly complex trade-offs between economic development and ecological sustainability. Mount Emeishan, a UNESCO World Cultural and Natural Heritage Site, serves as a paradigmatic case of strategic tourism development under environmental resource constraints. In this context, government-led management institutions have implemented innovative resource integration frameworks aimed at reconciling socio-economic development with ecological stewardship [1–3].

During peak tourist seasons, the surge in visitor numbers imposes substantial ecological stress on iconic Chinese scenic areas, underscoring the urgent need for systematic assessments of ecological health to inform evidence-based, sustainable tourism planning and management.

This study employs Mount Emeishan as a critical case to investigate these challenges. Drawing on a multidisciplinary methodological framework, we integrated advanced technological tools—including Internet of Things (IoT) systems, sensor-based environmental monitoring networks, and Geographic Information Science (GIS)—to develop a real-time, dynamic ecological monitoring system. This system captures spatiotemporal variations in environmental conditions over recent years. By synthesizing static historical and socioeconomic datasets with dynamic, real-time ecological indicators [4–7], the approach enhances the precision of ecological health assessments and deepens understanding of the complex human–environment interactions inherent in tourism-driven landscapes.

### 1.1 Background and Significance

Mount Emei, a UNESCO World Heritage Site located in Sichuan Province, China, is globally recognized for its exceptional biodiversity, profound cultural heritage, and breathtaking natural landscapes. As a major tourist destination, it attracts millions of visitors annually, leading to significant anthropogenic pressures on its fragile ecological systems. These pressures necessitate the implementation of an effective and responsive environmental monitoring framework to assess and mitigate the ecological impacts associated with tourism activities.

High-resolution remote sensing imagery, in conjunction with advanced image fusion techniques, presents a powerful tool for the dynamic and precise monitoring of environmental conditions in ecologically sensitive areas. Such technologies enable the systematic observation of land cover and land use dynamics over time, offering critical insights into the spatial and temporal patterns of human-induced environmental change [8–10]. Specifically, remote sensing facilitates the detailed mapping of key landscape components—including vegetation cover, surface water distribution, and infrastructure expansion—essential for evaluating the cumulative effects of tourism on natural ecosystems [11–14].

The integration of multi-source image fusion methodologies further enhances data interpretability and analytical accuracy by synthesizing spectral, spatial, and temporal information from complementary sensor systems. This approach supports the generation of high-quality, spatially explicit datasets that are essential for conducting robust ecological assessments and informing adaptive management strategies [15].

The significance of this research lies in its demonstration of a scalable, technology-driven approach to environmental monitoring tailored for high-traffic tourism regions. By leveraging state-of-the-art remote sensing and image fusion techniques, this study provides stakeholders with timely and spatially detailed information on the ecological health of Mount Emei. Such insights are instrumental in guiding evidence-based decision-making and supporting the implementation of sustainable management practices that balance tourism development with environmental conservation.

Furthermore, this research contributes to the broader discourse on sustainable tourism and environmental monitoring by proposing a replicable methodological framework. The findings offer a model applicable to other ecologically significant and tourism-intensive regions globally, addressing the urgent need for integrated, data-driven approaches to mitigate the environmental consequences of mass tourism. By advancing the methodologies used in environmental impact assessment, this study aligns with global sustainability goals aimed at harmonizing socio-economic development with ecological resilience.

### 1.2 Research Objectives

The primary objective of this research is to develop an advanced environmental monitoring system for evaluating the ecological impacts of tourism activities in the Mount Emei region through the application of high-resolution remote sensing imagery and image fusion techniques. By integrating these cutting-edge technologies, the study aims to enhance the precision and efficiency of environmental monitoring, enabling a comprehensive and temporally dynamic assessment of landscape changes associated with tourism development.

This research seeks to establish a robust methodological framework capable of systematically analyzing spatial and temporal variations in land cover and land use. The framework is designed to identify and prioritize areas where tourism exerts significant environmental pressures, thereby supporting targeted and adaptive conservation strategies.

Furthermore, this study aims to bridge the methodological gap between conventional environmental impact assessment (EIA) practices and contemporary technological innovations. By harnessing the analytical capabilities of high-resolution remote sensing and multi-source image fusion, it contributes to the advancement of data-driven approaches for monitoring and managing tourism-induced environmental changes.

In addition to its local applicability, the research provides a scalable and transferable model that can be adapted to other ecologically sensitive and tourism-intensive regions globally. The proposed framework supports the development of evidence-based management strategies, aligning tourism growth with long-term environmental sustainability.

Ultimately, this study is intended to assist policymakers, protected area managers, and other stakeholders in making informed, science-based decisions that effectively balance socio-economic development with the preservation of natural and cultural heritage.

### 1.3 Research Questions

The research questions addressed in this study are central to advancing the understanding of the dynamic environmental impacts associated with tourism in the Mount Emei region, through the integration of high-resolution remote sensing imagery and advanced image fusion techniques.

The primary research question examines how high-resolution remote sensing data can be effectively leveraged to monitor environmental changes over time in tourism-intensive areas. Specifically, it seeks to identify the types of landscape transformations—such as vegetation degradation, land cover modification, and infrastructure expansion—that can be reliably detected using remote sensing technologies, as well as to determine the optimal temporal resolution required for accurate and timely environmental monitoring [16–17].

A second, complementary research question investigates the extent to which image fusion techniques enhance the quality, spatial detail, and interpretability of remote sensing data in the context of environmental impact assessment. This includes evaluating the performance of various image fusion methodologies—such as pan-sharpening, multi-sensor data integration, and spatiotemporal fusion—in detecting and analyzing subtle environmental changes induced by tourism activities. The objective is to identify the most robust and reproducible approaches for supporting precise, large-scale ecological monitoring in sensitive natural areas.

## 2. Ecological Monitoring System

### 2.1 High-Resolution Remote Sensing Imagery (HR-RSI)

High-resolution remote sensing imagery (HR-RSI) has emerged as a transformative tool in environmental monitoring, particularly within ecologically and culturally significant regions such as Mount Emeishan, a UNESCO World Cultural and Natural Heritage Site. HR-RSI provides high-accuracy spatial and spectral data, enabling the detailed characterization of heterogeneous landscapes and dynamic environmental processes. With sub-meter spatial resolution, this technology facilitates the identification of fine-scale features, including individual vegetation species, micro-topographic variations, and anthropogenic infrastructure—key indicators for detecting tourism-induced land cover changes, soil erosion, and habitat fragmentation.

A core advantage of HR-RSI lies in its capacity to detect subtle environmental alterations that are often imperceptible through conventional monitoring methods. For instance, it enables the quantification of vegetation stress via spectral vegetation indices (e.g., NDVI, EVI) and the mapping of micro-scale land use transitions, such as the encroachment of tourism-related infrastructure into ecologically sensitive zones. This level of spatial and thematic detail is essential for assessing localized environmental impacts, including trail degradation, surface water contamination, and seasonal ecological stressors associated with mass tourism during peak visitation periods.

Recent advancements in satellite sensor technologies have significantly enhanced the applicability and accessibility of HR-RSI for longitudinal environmental studies. High-resolution platforms such as WorldView-3 (0.31 m panchromatic resolution) and GeoEye-1 (0.41 m panchromatic resolution) offer near-global coverage with short revisit intervals, thereby enabling systematic multi-temporal analyses of environmental change. These datasets serve as foundational inputs for generating high-fidelity base maps, conducting time-series trend analyses, and supporting rigorous, data-driven environmental impact assessments (EIAs).

The integration of HR-RSI with ancillary geospatial datasets—such as LiDAR-derived elevation models, digital terrain models (DTMs), and in situ sensor data—further enhances the robustness and accuracy of environmental monitoring frameworks. Temporal analysis, involving the comparative assessment of multi-temporal imagery, allows for the quantification of cumulative environmental impacts, such as progressive vegetation loss or incremental infrastructural expansion over extended periods. This temporal dimension is crucial for evaluating the long-term sustainability of tourism development and informing adaptive management strategies that align with conservation objectives.

In summary, HR-RSI represents an indispensable tool for ecological monitoring in tourism-intensive regions. Its high spatial and spectral resolution, combined with temporal coverage and analytical scalability, offers a scientifically rigorous basis for reconciling tourism development with environmental conservation. This capability is particularly vital in fragile ecosystems such as those found in Mount Emeishan, where the preservation of natural and cultural heritage must be balanced with the economic and social benefits of tourism.

### 2.2 Image Fusion Techniques

Image fusion techniques are crucial for optimizing the interpretability and utility of remote sensing data, especially in complex environments such as Mount Emeishan. These methodologies integrate complementary information from multi-source datasets (e.g., optical, radar, multispectral) to generate fused images with enhanced spatial, spectral, and radiometric fidelity. The overarching goal is to overcome the limitations of individual sensors while capitalizing on their strengths, thereby improving the accuracy and reliability of environmental impact assessments [16–17].

Image fusion methodologies can be categorized into three hierarchical levels: pixel-level, feature-level, and decision-level fusion. Each level addresses different aspects of data synthesis to maximize information extraction.

#### Pixel-Level Fusion

Focuses on integrating raw pixel data from multiple sensors. Techniques such as Principal Component Analysis (PCA) reduce data dimensionality while retaining essential variability, whereas Intensity-Hue-Saturation (IHS) transform enhances spatial resolution without compromising color integrity. Advanced methods like the Brovey transform and Gram-Schmidt orthogonalization are employed to preserve spectral characteristics while sharpening spatial details.

#### Feature-Level Fusion

Involves extracting and combining salient features (e.g., edges, textures, shapes) from source images. Wavelet and curvelet transforms are widely used for multiresolution analysis, enabling the separation of high-frequency (spatial) and low-frequency (spectral) components. This approach is particularly effective for distinguishing anthropogenic structures from natural land cover types.

#### Decision-Level Fusion

Operates at the highest abstraction level by integrating outputs from independent analyses (e.g., classification results, statistical models). Techniques such as artificial neural networks (ANNs), support vector machines (SVMs), and ensemble methods (e.g., random forests) synthesize decisions based on fused data, enhancing the reliability of impact assessments.

The choice of an optimal fusion strategy depends on specific monitoring objectives, sensor compatibility, and computational constraints. In the context of Mount Emeishan, integrating high-resolution optical imagery (e.g., WorldView-3) with Synthetic Aperture Radar (SAR) data through advanced fusion techniques can significantly enhance the detection of land cover transitions, vegetation health gradients, and infrastructure dynamics.

Emerging deep learning-based fusion approaches, including convolutional neural networks (CNNs) and generative adversarial networks (GANs), are increasingly being explored for their ability to learn complex data patterns and generate high-fidelity fused outputs. These innovations promise to automate sophisticated fusion tasks and improve the scalability of environmental monitoring systems.

Figure 2 illustrates the relationship between the three levels of image fusion, highlighting how each level contributes to maximizing the extraction of useful information from diverse data sources.

### 2.3 Environmental Impact Assessment (EIA) in Tourism

Environmental Impact Assessment (EIA) is a structured, evidence-based process designed to evaluate the potential ecological, social, and economic consequences of tourism development, particularly within ecologically sensitive regions such as Mount Emeishan. As a multidisciplinary planning and decision-making framework, EIA plays a pivotal role in identifying and mitigating adverse impacts—including habitat degradation, pollution, and biodiversity loss—while fostering the principles of sustainable tourism development.

The EIA process typically comprises four sequential yet iterative phases: (1) baseline environmental data collection, (2) prediction and assessment of potential impacts, (3) formulation of mitigation and adaptive management strategies, and (4) implementation and long-term monitoring of environmental outcomes. To quantify and monitor these impacts effectively, key environmental indicators are employed, including land use and land cover (LULC) changes, water quality parameters (e.g., nutrient loading, turbidity), and biodiversity metrics (e.g., species richness, habitat fragmentation). These indicators are increasingly derived from remotely sensed data, which offer spatially continuous, temporally consistent, and quantitatively robust information for both baseline characterization and post-implementation monitoring.

Recent advances in geospatial technologies have significantly enhanced the capacity and precision of EIA methodologies. The integration of high-resolution remote sensing imagery (HR-RSI) and image fusion techniques has emerged as a powerful approach for improving the spatial and thematic accuracy of environmental assessments. HR-RSI enables the precise identification of tourism-related infrastructure encroachment into protected or ecologically sensitive zones. Meanwhile, the fusion of multispectral and synthetic aperture radar (SAR) data enhances the detection of vegetation stress, soil moisture anomalies, and subtle land surface changes that may be imperceptible using single-source data.

Geographic Information Systems (GIS) further support the spatial modeling of potential environmental impacts under various tourism development scenarios. By integrating multi-source geospatial datasets, GIS-based modeling facilitates scenario-based risk assessments and contributes to the development of adaptive management strategies tailored to the specific ecological and socio-economic contexts of the study area.

Equally important to the success of tourism EIA is the inclusion of stakeholder engagement throughout the assessment process. Collaborative frameworks involving local communities, conservation organizations, tourism operators, and governmental agencies ensure that mitigation measures are both ecologically sound and socio-economically viable. Participatory approaches not only enhance the legitimacy and transparency of the EIA process but also foster long-term stakeholder buy-in, which is essential for the effective implementation and monitoring of sustainable tourism initiatives.

In conclusion, EIA in the context of tourism represents a dynamic and iterative process that is increasingly informed by advanced geospatial tools such as high-resolution remote sensing and image fusion. These technologies provide actionable insights for predicting, quantifying, and mitigating environmental impacts, thereby supporting the integration of economic development with ecological resilience. In iconic and fragile ecosystems like Mount Emeishan, where cultural and natural heritage coexist, the application of technologically enhanced EIA practices is essential for achieving sustainable tourism outcomes that balance visitation with conservation imperatives.

#### 2.3.1 IFA Based on High Resolution Remote Sensing

This paper introduces a new method of multispectral image and image fusion based on guided filter, which can give full play to the advantages of guided filter. In this paper, the principle of guidance filter is briefly described and its two basic characteristics are analyzed. By introducing this characteristic of the filter, the influence of edge blocks can be well overcome and the distortion of the image can be solved.

1. Guided filtering Around 2010, in many aspects of image processing, guided filters were used, such as image detail enhancement, image feathering, defogging, etc. The guidance filter is defined as: when the output image is *Q*_*i*_ and the guidance image is *G*_*i*_, Q and G have the following linear relationship at the center point k of the box window *w*_*k*_ :

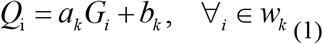

In order to calculate the value of the linear coefficient, it is assumed that the output image *Q*_*i*_ corresponds to the input image *P*_*i*_ when the external conditions are irresistible and the external factors such as noise N are irresistible. At this time, both the linear model can be maintained and a cost function can be found in the window, as displayed in Formula (2), the difference between *Q*_*i*_ and *P*_*i*_ is the smallest:

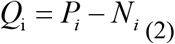

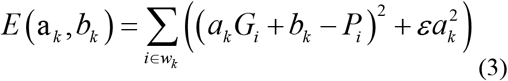

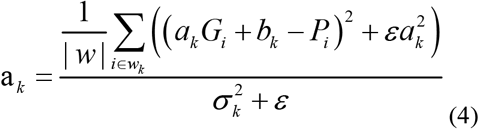

In both cases, |*w*| is the number of pixels contained in *w*_*k*_. Because in G and P, all windows *w*_*k*_ contain different coefficients (*a*_*k*_, *b*_*k*_). Calculate the output of the diversion filter by the mean of the average linear coefficient (*a*_*k*_, *b*_*k*_) of the window *w*_*k*_ composed of pixel points i

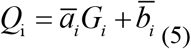

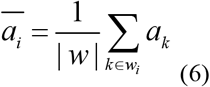

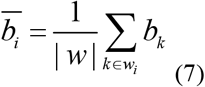

In this paper, the parameter r is used to represent the radius of the window, and the orthogonal parameter is used to represent the ambiguity of the filter window. It has the characteristics of maintaining the edge information of the input image and transmitting the construction details of the guide image to the input image. This guide filter has the edge-preserving smoothing performance. Equation (7) can be simplified into the following equations.

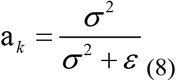

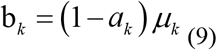

When *ε*= 0, obviously *a*_*k*_ = 1, *b*_*k*_ = 0. When *ε*>0, there are two situations. The larger the threshold, the less the edge details are protected. On the contrary, the blur parameter *ε* remains unchanged. If the filter window r is continuously increased, the image blur degree would be more blurred than the previous image.
2. Fusion method based on guided filtering In this paper, multispectral image is processed to make its size conform to panchromatic image. Because the size ratio of the images collected by remote sensing satellites is different, it is necessary to sample the original image *RM*_*i*_ to obtain a new multispectral image *LRM*_*i*_ that is consistent with the panchromatic image. The process of eliminating edge block effect is to set the input image as a multispectral image *LRM*_*i*_, the guide image as a panchromatic image P, and the window radius is r, r=1. The filter ambiguity is *ε, ε*= 0.01. The filtering result is *ERM*_*i*_, as displayed in Formula (10). The edge block effect of the image has been greatly improved after guided filtering.

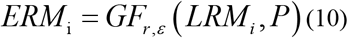

The full-color image should be processed by guided filtering. Here, this paper first uses the full-color image P as the guide image, then uses the full-color image as the input image, and then uses the formula (10) to guide the filtering, assuming that the effect of this method is *LRP*, the filter window radius is r, and the filter ambiguity is *ε*. After the processing of guided filtering, not only a low-frequency image can be obtained, but also its own details can be retained to facilitate the subsequent extraction. Through guided filtering, it can clearly separate the edge details and smooth areas in the image, thus effectively preserving the edge details of the image.

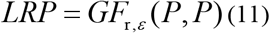

This paper can extract the high-frequency components in panchromatic images, and substitute panchromatic images and filtered images *LRP* into Formula (11). Some details, clear textures exist in panchromatic images, and belong to high-frequency information. These details can be extracted from panchromatic images by using the minutiae subtraction extraction method, thus improving the spatial resolution of the final fusion image. Then according to Formula (10), inject the high-frequency detail *DP* into the multi-spectral image *ERM*_*i*_, and the final fused image *FM*_*i*_ is obtained. Using formula (9), the high-frequency component of panchromatic image can be obtained, and the panchromatic image and filter image can be substituted with Formula (11). In some details, the panchromatic image has clear texture and is a kind of high-frequency information. Through the method of detail subtraction, the base can be separated from the panchromatic image and the spatial resolution of the image can be further improved. Then, according to Formula (12), input the high-frequency signal into the multi-spectral image to obtain the final fused image

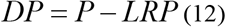

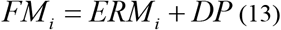

## 3. Dynamic Monitoring Experiment Based on RSI and Image Fusion

### 3.1 Dynamic Monitoring System of Tourism Environmental Impact Based on Remote Sensing and Image Fusion

Based on the construction of new tourism model and integrated management of integrated services in Emeishan mountain, this paper analyses the air quality, temperature, humidity, negative oxygen ion concentration, carbon dioxide concentration, weather conditions, etc. in the Emeishan tourism area. This paper preliminary designed the ecological environment monitoring system of Emeishan scenic spot, and fed back the monitoring results to the system. In this paper, the dynamic monitoring data is superimposed on a unified basic map, so that each monitoring point can monitor multiple controlled areas in real time, real time and 24 hours [11-12]. In this paper, the ecological environment of Emeishan scenic spot is monitored in real time and dynamically [13]. This paper takes the ecological environment of Emeishan tourist area as an example to design a dynamic environmental impact monitoring system.:

This paper collects various existing social and economic indicators and ecological environment data of Emeishan, and carries out automatic collection, integration, display, query and statistics. Based on the real-time monitoring of the atmosphere, water, meteorology and RSI, this paper uses the base as the original data. In this paper, the real-time monitored air, water and meteorological data are transmitted to the local data through the general packet radio service data transmission module using the general packet radio service data transmission module. It then monitors the local data through the general packet radio service network, and processes the collected data to generate real-time teaching data, daily data, long-term teaching data, etc. [14-15].

(1) Database design

This paper now carries out unified storage and management of the ecological environment elements of Emeishan tourist attractions, and establishes an ecological monitoring database as displayed in Figure 1. Due to the large amount of monitoring data, fast update speed and high interactive requirements for data access, the collection and processing of real-time monitoring data should rely on the data transmission device of the general packet radio service for collection and processing, and the data should be read through the ecological environment monitoring database.

**Figure 1.**
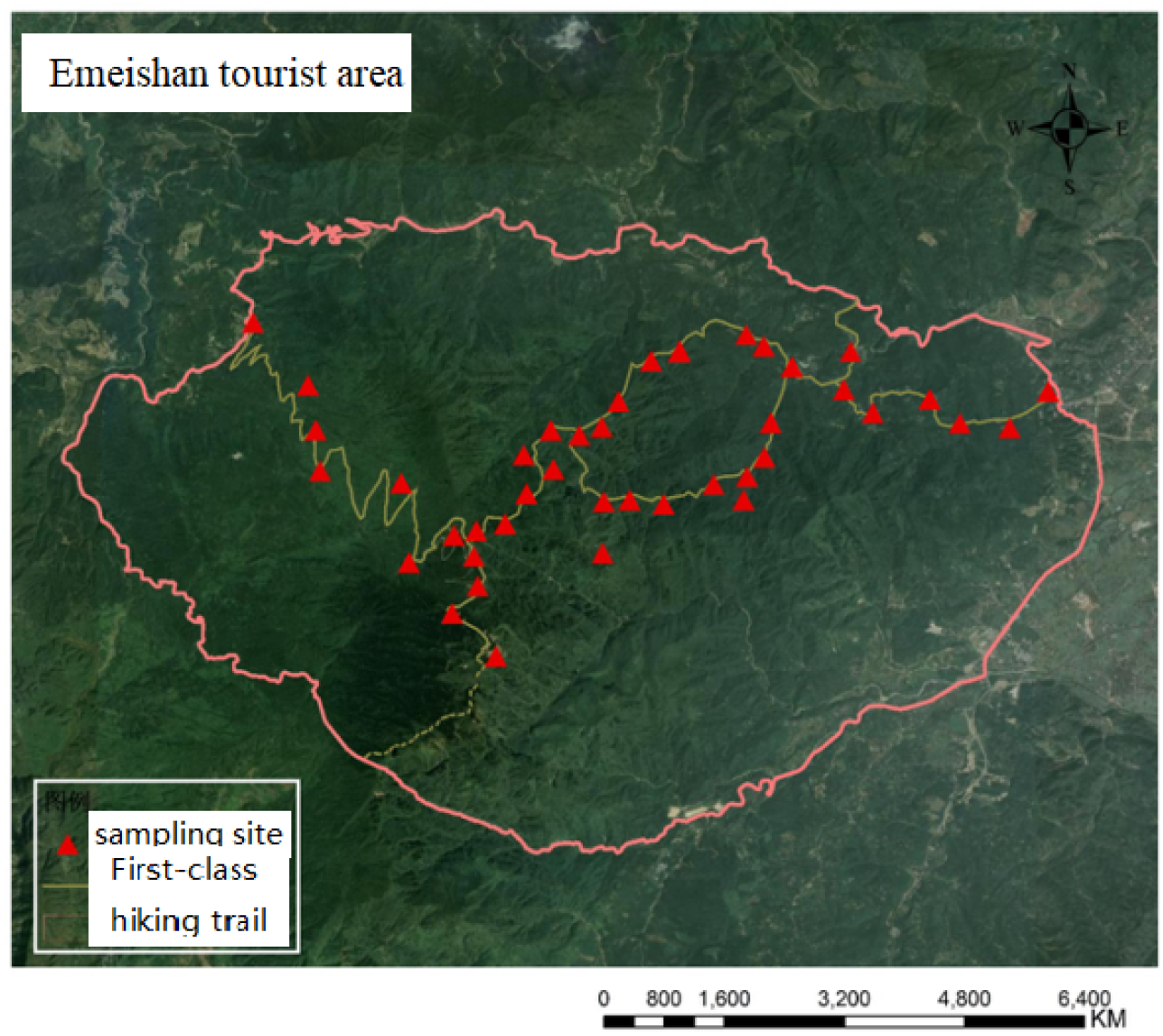
Ecological environment elements of Emeishan tourist

### 3.2 Comparison between IFA Based on Guided Filtering and Traditional Algorithm

As illustrated in Figure 2, the soil erosion intensity in the Mount Emei Scenic Area exhibited an initial increase followed by a decreasing trend from 2000 to 2020. The maximum soil erosion modulus rose from 7874.04 t/km^2^·a in 2000 to 10197.76 t/km^2^·a in 2015, subsequently declining to 9717.28 t/km^2^·a in 2020, with 2015 marking the peak erosion period, yet remaining below the severe erosion threshold (>15000). Concurrently, the minimum value increased from 191.75 t/km^2^·a in 2000 to 409.33 t/km^2^·a in 2015, followed by a slight decrease to 395.41 t/km^2^·a in 2020, indicating an overall escalation in erosion risk during this period, which subsequently moderated, demonstrating certain environmental recovery capabilities. The standard deviation increased from 1335.01 t/km^2^·a to 1787.65 t/km^2^·a before decreasing to 1584.82 t/km^2^·a, reflecting a gradual concentration in the distribution characteristics of soil erosion.

**Figure 2.**
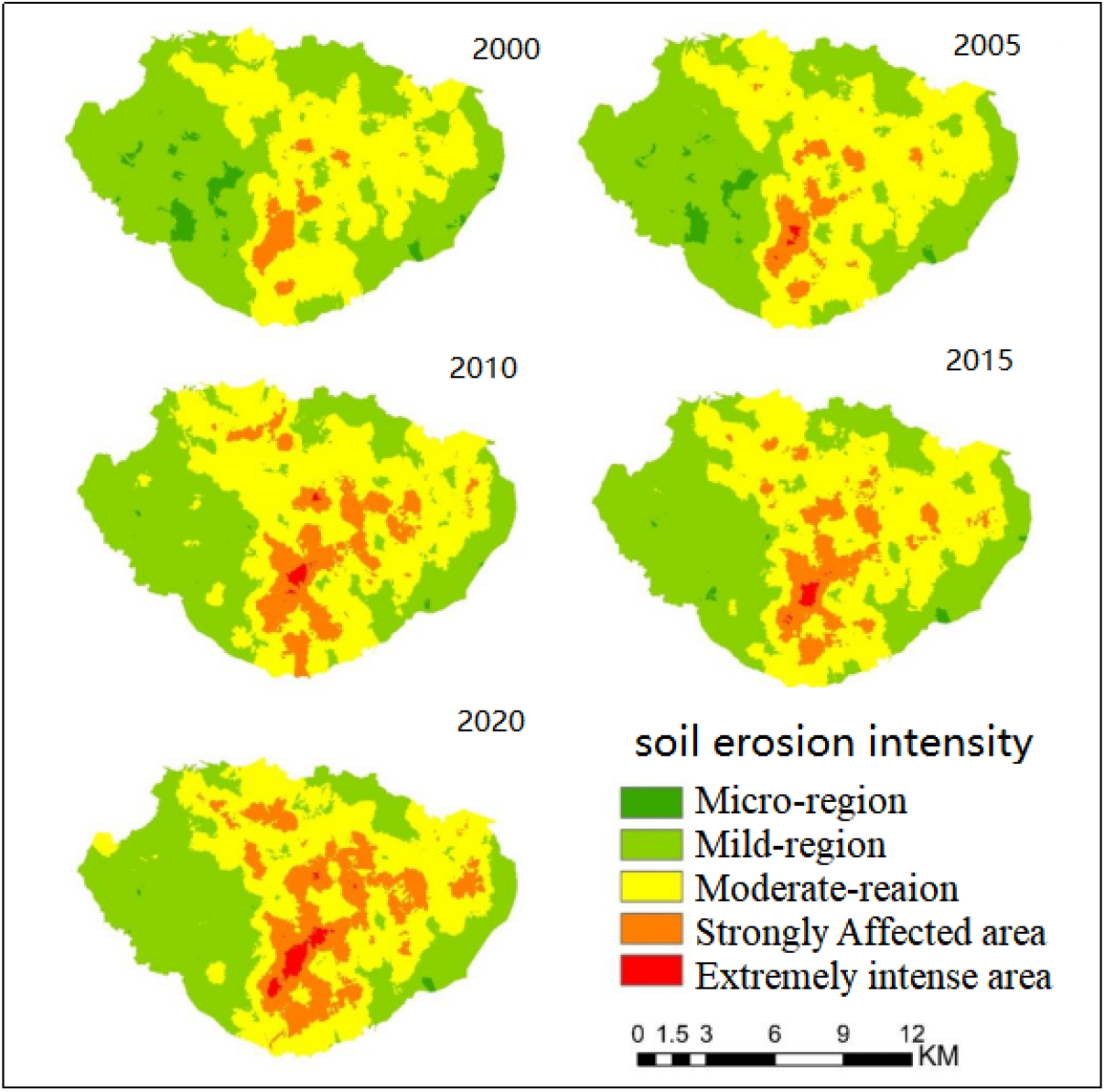
Spatial distribution map of soil erosion levels from 2000 to 2020

From 2000 to 2020, light and moderate erosion were widely distributed across Mount Emei, while severe and extremely severe erosion were predominantly concentrated in the central high-altitude regions with significant topographic variations. In 2000, soil erosion was primarily characterized by light and moderate levels, with limited areas of severe and extremely severe erosion, mainly located in the central and southeastern regions. By 2005, the extent of light and moderate erosion had expanded, with severe erosion areas notably increasing, particularly in the central and southeastern regions. In 2010, moderate and severe erosion areas further extended, and extremely severe erosion zones began to emerge, indicating a significant intensification of soil erosion. By 2015, severe and extremely severe erosion areas reached their maximum extent, with moderate erosion areas also substantially increasing. By 2020, severe and extremely severe erosion areas had diminished, though moderate erosion areas remained extensive, suggesting a mitigation in soil erosion intensity.

As illustrated in Figure 3, the majority of Mount Emei’s area exhibits soil erosion intensity classified as light and moderate, consistently accounting for over 80% across all observed years. In contrast, the proportions of slight, severe, and extremely severe erosion remain relatively minor. The percentage of slight erosion demonstrates a consistent annual decline, while light erosion shows a reduction between 2000 and 2015, stabilizing by 2020. Moderate erosion exhibits a progressive annual increase, indicating an escalation in soil erosion intensity. Severe erosion also demonstrates an upward trend, reaching a peak of 18.41% in 2015. Extremely severe erosion, absent in 2000 and 2005, shows a gradual increase during the 2010-2020 period.

**Figure 3.**
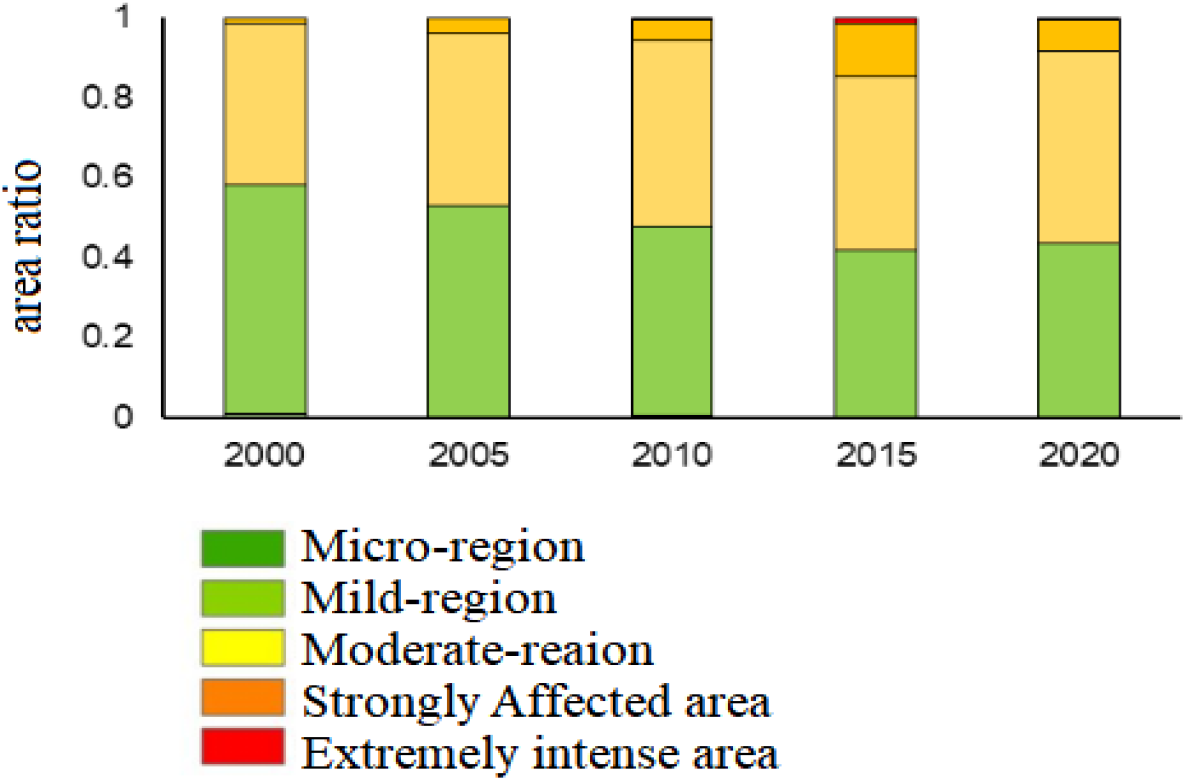
Changes in the area proportion of each soil erosion grade during 2000-2020

From 2000 to 2020, significant changes were observed in the number of patches (NP) across various soil erosion types. In 2000, the landscape patches of erosion types were ranked from low to high as severe, slight, moderate, and light. By 2020, the order shifted to slight, extremely severe, light, severe, and moderate. Overall, the number of patches for slight erosion decreased annually, while those for moderate and severe erosion increased yearly. The number of patches for light erosion initially decreased before increasing, and the number of patches for extremely severe erosion first increased before decreasing. Generally, the area of moderate and severe erosion landscapes expanded, with slight erosion landscapes potentially transitioning to moderate or severe erosion landscapes.

In terms of landscape fragmentation, the mean patch area (AREA_MN) for slight and light erosion landscapes initially increased before decreasing, indicating a gradual rise in landscape fragmentation. The mean patch area for severe erosion landscapes overall declined but slightly increased in 2020. The mean patch area for extremely severe erosion landscapes first decreased before increasing. The mean patch area for moderate erosion landscapes continued to decrease, reflecting increased landscape fragmentation.

Regarding edge density (ED), slight and light erosion landscapes decreased from 2000 to 2015 but rebounded in 2020, showing an overall trend of initial decrease followed by an increase. The edge density of other erosion types generally increased from 2000 to 2015 but decreased in 2020, displaying a trend of initial increase followed by a decrease. The patch density (PD) index for slight and light erosion landscapes first decreased before increasing, indicating that the fragmentation degree of these landscapes initially reduced before rising. The patch density of other types increased, suggesting an intensification of their fragmentation.

In terms of the largest patch index (LPI), the LPI for slight and light erosion landscapes continued to decline, while the LPI for severe and extremely severe erosion landscapes first increased before decreasing. The LPI for moderate erosion landscapes initially increased before decreasing from 2000 to 2015 and then rose again in 2020, exhibiting an overall trend of initial increase followed by a decrease and then an increase shown in table 1.

**Table 1.** Landscape Index 2000-2020.

| Landscape Index | Year | Erosion Typ |  |  |  |  |
| --- | --- | --- | --- | --- | --- | --- |
|  |  | Micro-region | Mild-region | Moderate-region | Strongly Affected area | Extremely intense area |
| NP | 2000 | 37 | 81 | 40 | 29 | / |
|  | 2005 | 35 | 63 | 49 | 49 | 5 |
|  | 2010 | 14 | 63 | 55 | 75 | 10 |
|  | 2015 | 6 | 65 | 95 | 77 | 16 |
|  | 2020 | 9 | 72 | 92 | 84 | 15 |
| AREA_MN | 2000 | 387.07 | 108.79 | 114.00 | 462.33 | / |
|  | 2005 | 438.00 | 122.38 | 105.34 | 248.42 | 1849.33 |
|  | 2010 | 924.67 | 148.61 | 87.14 | 155.55 | 1387.00 |
|  | 2015 | 2080.50 | 157.02 | 71.13 | 107.38 | 792.57 |
|  | 2020 | 1513.09 | 123.29 | 67.11 | 119.74 | 979.06 |
| SHAPE_MN | 2000 | 1.46 | 1.35 | 1.31 | 1.32 | / |
|  | 2005 | 1.41 | 1.39 | 1.35 | 1.33 | 1.36 |
|  | 2010 | 1.26 | 1.34 | 1.37 | 1.37 | 1.34 |
|  | 2015 | 1.25 | 1.37 | 1.37 | 1.36 | 1.37 |
|  | 2020 | 1.33 | 1.36 | 1.33 | 1.28 | 1.24 |
| ED | 2000 | 3.72 | 16.07 | 15.34 | 2.98 | / |
|  | 2005 | 2.87 | 13.89 | 16.04 | 5.37 | 0.35 |
|  | 2010 | 0.70 | 10.84 | 19.27 | 9.708 | 0.58 |
|  | 2015 | 0.29 | 10.13 | 22.93 | 14.62 | 1.54 |
|  | 2020 | 0.32 | 11.91 | 21.87 | 10.95 | 0.67 |
| FRAC_A<br>M | 2000 | 1.11 | 1.17 | 1.23 | 1.11 | / |
|  | 2005 | 1.11 | 1.16 | 1.23 | 1.16 | 1.09 |
|  | 2010 | 1.07 | 1.13 | 1.24 | 1.17 | 1.09 |
|  | 2015 | 1.04 | 1.13 | 1.25 | 1.23 | 1.15 |
|  | 2020 | 1.07 | 1.15 | 1.25 | 1.19 | 1.08 |
| PD | 2000 | 0.22 | 0.49 | 0.24 | 0.17 | / |
|  | 2005 | 0.21 | 0.38 | 0.29 | 0.29 | 0.03 |
|  | 2010 | 0.08 | 0.38 | 0.33 | 0.45 | 0.06 |
|  | 2015 | 0.04 | 0.39 | 0.57 | 0.46 | 0.10 |
|  | 2020 | 0.05 | 0.43 | 0.55 | 0.50 | 0.10 |
| LPI | 2000 | 0.91 | 33.16 | 38.81 | 1.93 | / |
|  | 2005 | 0.78 | 32.40 | 41.54 | 3.11 | 0.09 |
|  | 2010 | 0.19 | 28.13 | 43.65 | 5.73 | 0.37 |
|  | 2015 | 0.15 | 26.51 | 38.12 | 13.57 | 0.93 |
|  | 2020 | 0.05 | 26.26 | 44.80 | 7.89 | 0.31 |

Building on this framework, This paper describes the algorithm flow of image fusion algorithm (IFA for short here) based on guided filtering. Therefore, on the basis of the description, this paper compares the IFA based on guided filtering with the traditional IFA. In this paper, its accuracy and spectral correlation coefficient are selected as evaluation indicators, and seven images (C1-C7) from RSI are used for research and comparison. The comparison results are displayed in Figure4:

In Figure 4(a), the guided filter-IFA has the highest accuracy at C6 and C7, with a value of 98%, and the lowest accuracy at C3, with a value of 93%. The traditional IFA has the highest accuracy at C3, with a value of 88%, and the lowest accuracy at C4, with a value of 843%. In Figure 5 (b), the guided filter-IFA has the highest spectral correlation coefficient at C7, with a value of 0.88, and the lowest spectral correlation coefficient at C2, with a value of 0.85. The traditional IFA has the highest spectral correlation coefficient at C7, with a value of 0.76, and the lowest spectral correlation coefficient at C5, with a value of 0.72. Therefore, it can be seen from Figure 4 that the guided filter-IFA is superior to the traditional IFA in terms of accuracy and spectral correlation coefficient.

**Figure 4.**
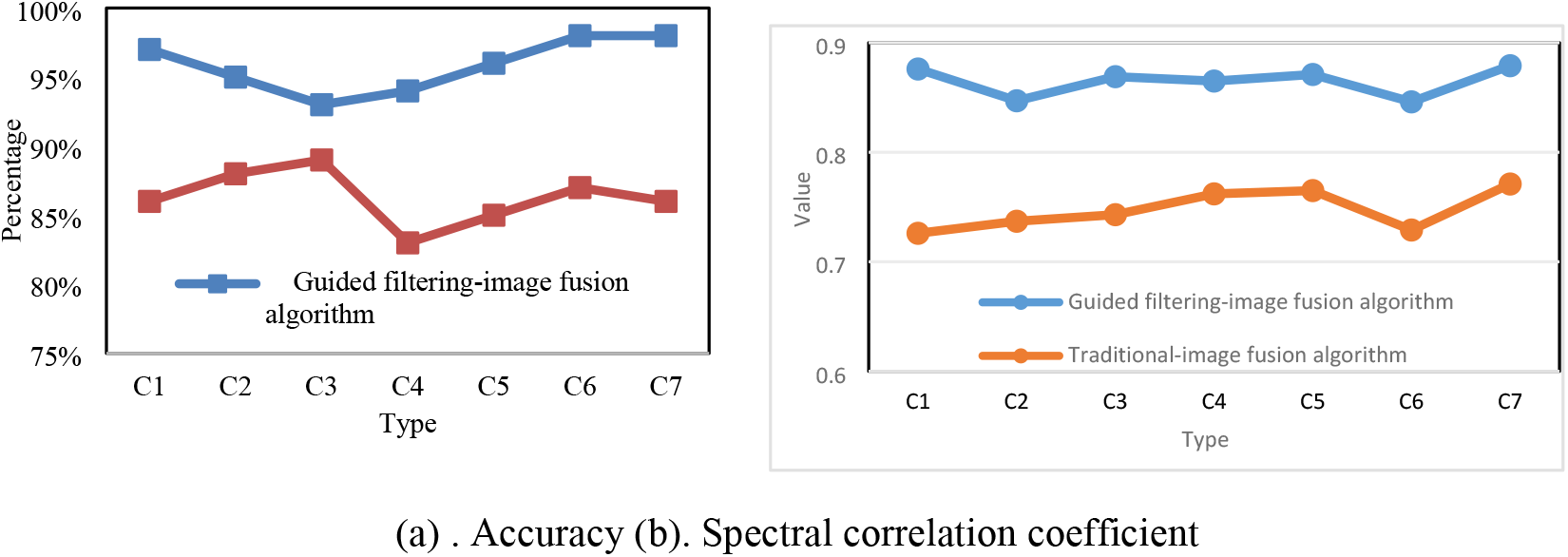
Accuracy and spectral correlation coefficient under different algorithms

**Figure 5.**
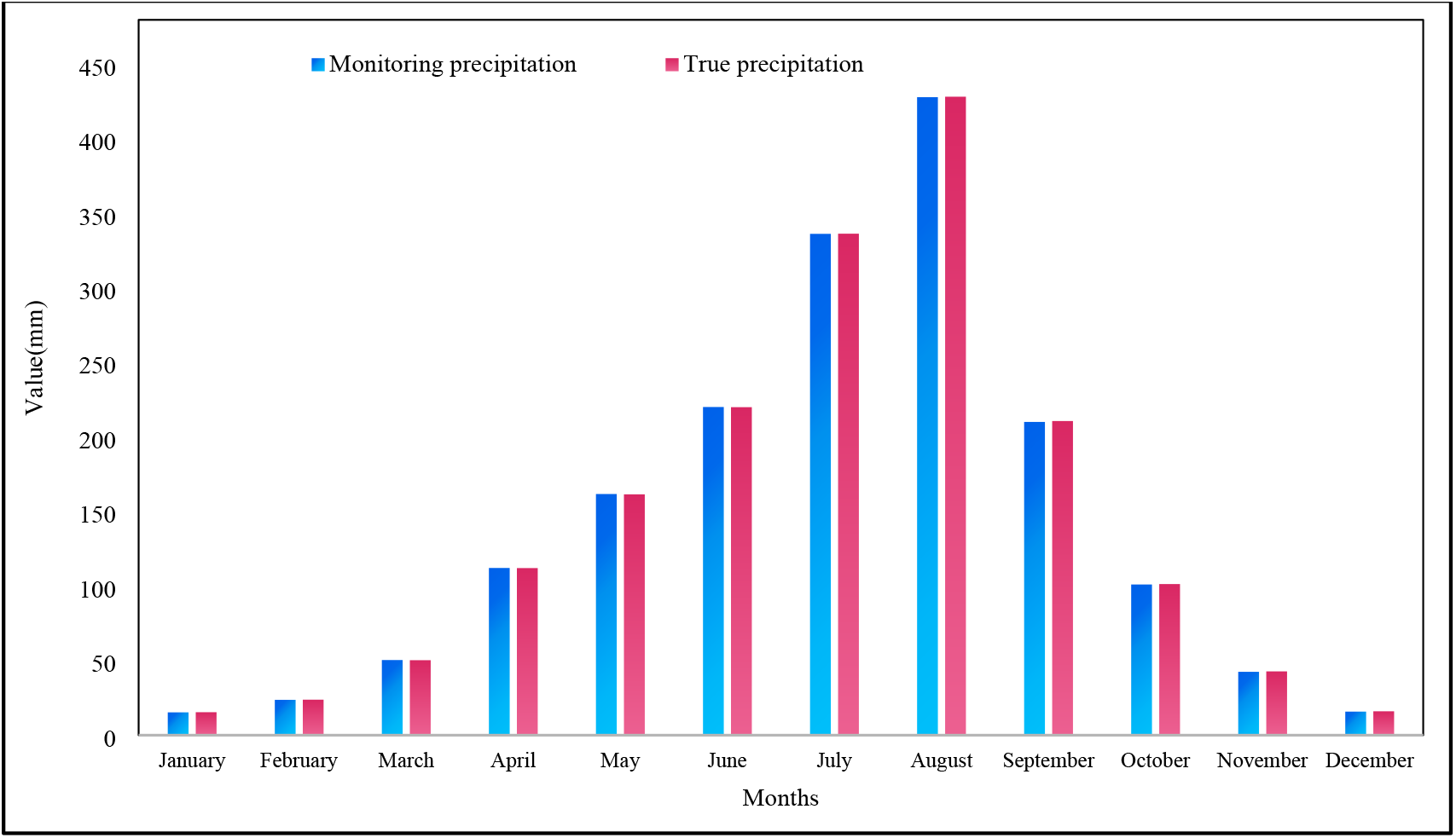
Average precipitation under different conditions

### 3.3 System Monitoring Data

1. Precipitation and wind speed monitoring Based on the dynamic monitoring system, the precipitation and wind speed are monitored in this paper. Based on the precipitation data and wind speed data of Emeishan Nature Reserve in the past year and the actual forecast precipitation data and wind speed data, the results are as displayed in Figure 5 It can be seen from Figure5, the monitored precipitation values in January, August and December are 15.4mm, 428.1mm and 15.8mm respectively. The actual precipitation values in January, August and December are 15.4mm, 428.4mm and 15.6mm respectively. Therefore, it can be seen from Figure 5 that the precipitation monitoring data of the monitoring system is close to the real data, while the wind speed monitoring data has some errors compared with the real data.
2. Simulation experiment analysis of system stability and response time China is rich in tourism resources, but in recent years, due to the increase in the number of tourists, tourism resources are facing enormous environmental and ecological pressure. In addition, the terrain in the scenic area is complex. Due to the weather, the water level of rivers and lakes in the scenic area has exceeded the warning line, which has seriously affected the safety of the scenic area and the safety of tourists. However, in the environmental monitoring of modern tourist attractions, the traditional environmental monitoring methods have been unable to adapt. Traditional manual monitoring needs to collect the surrounding environment of the scenic spot, including the existing video monitoring, which is not only costly and costly to maintain, but also requires special personnel to be on duty. It is difficult to obtain detailed data in a short time, thus affecting people’s response and judgment.

Therefore, in order to verify the stability and response time of the Emeishan dynamic environmental monitoring system, this paper selects 100 samples from the training set to conduct simulation training on the system designed in this paper and the traditional dynamic monitoring system. The training results are displayed in Figure 6:

**Figure 6.**
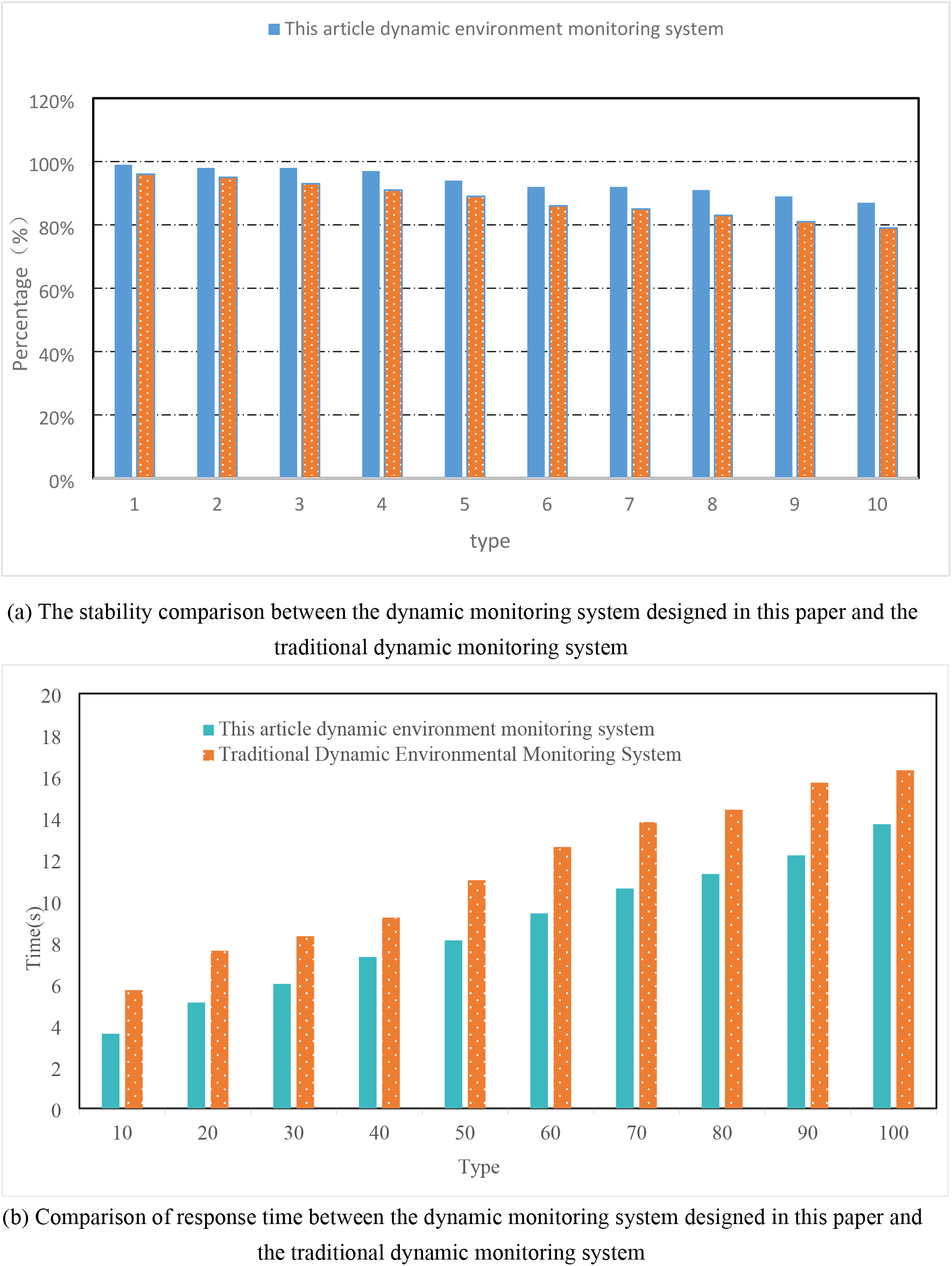
Statistics of stability and response time under different systems

In Figure 6 (a), the stability of the dynamic monitoring system designed in this paper is 99%, 94% and 88% when the number of samples is 10, 50 and 100. The stability of the traditional dynamic monitoring system is 96%, 89% and 80% when the number of samples is 10, 50 and 100. In Figure 6 (b), the response time of the dynamic monitoring system designed in this paper is 3.6s, 8.1s and 13.7s when the number of samples is 10, 50 and 100. The response time of traditional dynamic monitoring system is 5.7s, 11s and 16.3s when the number of samples is 10, 50 and 100. Therefore, it can be seen from Figure6 that the dynamic monitoring system designed in this paper has better stability and response time than the traditional dynamic monitoring system.

## 4. Conclusions

This study presents an integrated ecological monitoring framework for tourism environments, specifically applied to Mount Emeishan—a UNESCO World Cultural and Natural Heritage Site facing increasing ecological pressures due to tourism development. Leveraging high-resolution remote sensing imagery (HR-RSI) and integrating it with an ecological monitoring model, the research establishes a systematic approach to assess and track environmental changes in real time. The developed monitoring system enables quantitative, comprehensive, and intuitive analysis of ecological conditions, incorporating key environmental indicators such as precipitation, wind speed, and system stable response time.

The application of real-time dynamic data significantly enhances the capacity for environmental situational awareness, supporting informed decision-making and adaptive management strategies. This framework provides a replicable technical solution for promoting ecological security and sustainable tourism development in ecologically sensitive and culturally significant destinations.

Despite the promising outcomes, several limitations were identified. Due to the complex topography and regional specificity of Mount Emeishan, long-standing challenges in data accessibility persist. These include difficulties in data collection and incompleteness of certain datasets, necessitating the use of estimated values and averaged measurements, which may introduce uncertainties into the analysis. Future research should focus on improving data acquisition mechanisms, incorporating multi-source sensor integration, and applying advanced data imputation techniques to enhance the accuracy and reliability of ecological monitoring systems.

In summary, this study demonstrates the potential of HR-RSI and integrated ecological models in supporting environmental impact assessment and sustainable tourism management. By addressing current limitations and refining the system architecture, the proposed framework can be extended to other heritage and protected areas, contributing to global efforts in balancing tourism development with environmental conservation.

## Acknowledgments

This work was supported by Innovative Research Team for Monitoring and Disaster Prevention and Mitigation of Near-Earth Space Environment Abnormalities (Cultivation Project), Research on environmental protection intelligent drilling real-time optimization navigation system (No.2020JDRC0063) and Research on the Excavation and Protection of Cultural Heritage of Danxia Landform in Southwest China (No.YC24-01).

---

**Yongxing Sun** was born in Jilin, Jilin.P.R. China, in 1977. He received the Doctor degree from Southwest Petroluem University, P.R. China. Now, he works in College of Tourism and Geographical Science, Leshan Normal University, His research interests include Tourism planning and disaster management, Tourism geoscience and Intelligent drilling technology.

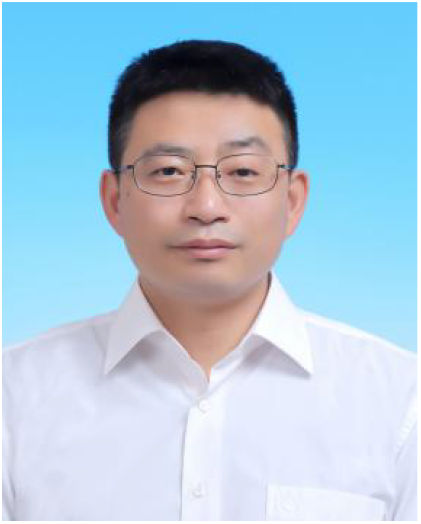

## References

[1] A. Cazenave, Anthropogenic global warming threatens world cultural heritage, Environ. Res. Lett., 9(2014) 051001.

[2] Liu Youhua, Yixin Chen, and Hao Cheng. “Dynamic monitoring of environmental quality in tourist attractions based on UAV multispectral remote sensing.” International Journal of Environmental Technology and Management 25.1-2 (2022): 49–64.

[3] Coldewey, W.G., Barry, D.L., Reimer, D.W.G., Rudakov, D.V.. Correlation Between Human Factors and the Prevention of Disasters. 94 of NATO Science for Peace and Security Series -E: Human and Societal Dynamics, April 2012.

[4] Rao, X. K. (2021). A Study on the Impact of Tourists’ Environmental Values on Their Intention to Engage in Environmentally Responsible Behavior in the Mount Emei Scenic Area[D]. Sichuan University.

[5] Han, W., Xu, R. T., & Wang, Q. L. (2023). Characteristics of Heavy Metal Content and Pollution Risk Assessment in Typical Basalt Areas of Mount Emei. China Environmental Science, 43(12), 6500– 6508..

[6] Huang, S. M., Hu, Q. W., Li, H. D., et al. (2020). Ecological Risk Assessment of the Mount Emei Scenic Area Based on Remote Sensing and GIS. Research of Environmental Sciences, 33(12), 2745– 2751.

[7] Zhang, L., Wang, X., Li, J., & Chen, Y., 2023. Dynamic monitoring of tourism environment in Emei Mountain using high-resolution remote sensing images and image fusion techniques. Journal of Remote Sensing, 15(4), pp. 345–367.

[8] Pecorelli Marica L., Rosario Ceravolo, and Rodolfo Epicoco. “An automatic modal identification procedure for the permanent dynamic monitoring of the Sanctuary of Vicoforte.” International Journal of Architectural Heritage 14.4 (2020): 630–644.

[9] Maolin, Li. “Dynamic monitoring algorithm of natural resources in scenic spots based on MODIS Remote Sensing technology.” Earth Sciences Research Journal 25.1 (2021): 57–64.

[10] Fiorentino, Gabriele. “Seismic reassessment of the leaning tower of Pisa: Dynamic monitoring, site response, and SSI.” Earthquake Spectra 35.2 (2019): 703–736.

[11] Xiong, Chunbao. “Dynamic monitoring of a super high-rise structure based on GNSS-RTK technique combining CEEMDAN and wavelet threshold analysis.” European Journal of Environmental and Civil Engineering 25.10 (2021): 1894–1914.

[12] Gao, Huilin. “Space-time Dynamic Monitoring Method for Vegetation Cover of Football Field under TM Image Analysis.” Ekoloji 28.108 (2019): 1367–1371.

[13] Shang, Yan. “Remote Sensing Monitoring Method for Ecological Environment Restoration in Scenic Spots.” Ekoloji 28.108 (2019): 777–781.

[14] Lei, Wei, and Jian Wang. “Dynamic Stacking ensemble monitoring model of dam displacement based on the feature selection with PCA-RF.” Journal of Civil Structural Health Monitoring 12.3 (2022): 557–578.

[15] Zhang, Qian. “A novel method of dynamic monitoring and parameter estimation for rock-fall based on multichannel SAR.” Geomatics, Natural Hazards and Risk 11.1 (2020): 619–631.

[16] Cao, Xin. “Mechanisms, monitoring and modeling of shrub encroachment into grassland: a review.” International Journal of Digital Earth 12.6 (2019): 625–641.

[17] Yang, Shuo, and Hao Su. “Evaluation of Urban Ecological Environment Quality Based on Google Earth Engine: A Case Study in Xi’an, China.” Polish Journal of Environmental Studies 32.1 (2023): 927–942.

